# MicroRNA 106b: Role in the reprograming of mitochondrial machinery and carcinogenesis in hepatic cells

**DOI:** 10.1101/2024.04.25.591197

**Authors:** Ashutosh Kumar Maurya, Lincy Edatt, V.B. Sameer Kumar

## Abstract

Cancer is a disease of unregulated cell growth. The process of initiation and progression of cancer is called carcinogenesis and the factors possessing ability to induce carcinogenesis are called carcinogens. Along with the coding sequence, the non-coding sequence also play very crucial role in the process of carcinogenesis. MicroRNAs are small non-coding RNAs having targets on both the classes of genes important in cancer i.e., oncogenes and tumour suppressor genes, thus act as key play in carcinogenesis. Dysfunctional mitochondrial metabolism has been widely reported in cancer and this malfunctioning could be brought in by suppression of the expression pattern of important mitochondrial genes by microRNAs. Our in-silico analysis revealed that miR 106b possess targets on several important mitochondrial genes involved in various complexes of electron transport chain. Further, we checked the role of miR 106b in reprogramming of the mitochondrial mechanism and carcinogenesis. The results suggested that miR 106b contributes to carcinogenesis in hepatic cells by modulating the mitochondrial metabolism.

## Introduction

One of the vital organelles for a cell’s survival and metabolism is the mitochondria, which are involved in the production of energy, required by the cells to perform various cellular actions (Liu Y et. al, 2023). Apart from their role in normal functioning of the cells, mitochondria play regulatory role in many human diseases ranging from inborn errors of metabolism, cardiovascular, alzheimer’s and cancer (Annesley S et. al, 2019, Swerdlow R, 2012, Cueva J et. al, 2002, Alam M et. al, 2016). These conditions progress primarily owing to the dysfunctional mitochondrial machinery (San-Millan I, 2023).

The mitochondrial function is regulated critically and can be determined by both nuclear and mitochondrial encoded genes (Kummer E et. al, 2021). The excess or down regulation of any gene involved in maintaining its normal physiological levels is undesirable for the normal physiology of mitochondria (Dorji J et. al, 2020). The RNA and protein levels in normal physiological as well as pathological conditions are regulated by one of the key regulators, small non-coding RNA molecules called as miRNA (Zhang P. et. al, 2019).

MicroRNAs are small non-coding RNAs (Saliminejad K et. al, 2019), which play important role in carcinogenesis by targeting genes involved in cell division and proliferation (Pavet V et. al, 2011) Apart from this, microRNAs could also regulate the expression pattern of the genes involved in mitochondrial functioning leading to reprogramming of the mitochondrial system of energy production (Suriya M et. al, 2022). The altered mitochondrial metabolism has been reported as the hall mark of a variety of cancer (Bosc C et. al, 2017).

In cancer, the mitochondrial machinery is rewired and glycolytic pathway is activated as alternate source of energy production (Schiliro C et. al, 2022, Shiratori R et. al, 2019). Micro RNAs plays an important role in this metabolic switch, as they target various genes involved in the mitochondrial metabolism, which could either be of nuclear or mitochondrial origin (Arora A et. al, 2015).

MicroRNA 106b is found on 7th chromosome at 7q22.1 location and it has been reported to be involved in various cancers by regulating the genes involved in cellular proliferation, invasion and metastasis (Enkhnaran B et. al, 2022, Yang C et. al, 2021, Yang F et. al, 2022). It has been found that miR 106B is highly expressed in various cancer including colon cancer, lung cancer, breast cancer etc. (Zhuang M et. al, 2019, Wang Z et. al, 2020, Li N et. al, 2017). This study deals with assessment of the role miR 106b in reprogramming of mitochondrial machinery and carcinogenesis.

## Methodology

### Cell culture

Hela (cervical cancer cell line), WRL-68 (Human hepatic non cancerous cell line) and HepG2 cells (Hepatic carcinoma cell line) were cultured in DMEM supplemented with 10% FBS, antibiotic-antimycotic solution and L-Glutamine. The cells were maintained under standard culture conditions at 37°C with 5% CO2 and 95% humidity. For experiments, seeding density of 0.4 × 10^4^ cells (96 well), 0.6×10^6^ cells (30mm dish), 0.8×10^6^ cells (60mmdish), 2.2×10^6^ cells (100 mm dish) were used.

### miR 106b cloning

miRNA 106b was cloned in pCMV miR vector between BamH1 and Xho1 restriction sites and successful cloning was confirmed by sequencing.

#### Transformation

The competent cells (DH5 α) were transformed with miR 106b plasmid by heat shock method where the plasmid was incubated with the competent cells followed by a quick heat shock at 90°C for 2 minutes and then immediately transferring it on ice. The transformed cells were then plated on agar plate containing kanamycin.

### Plasmid isolation

A Single colony was picked from the agar plate and grown in the LB broth containing Kanamycin. The broth was incubated at 37 degree C in a shaking incubator. The plasmid was isolated by using Himedia midi kit following manufacturers protocol.

### Transfection

HeLa cells were seeded in 6 well plates and grown in a monolayer. After reaching 70% confluency, the cells were transfected with miR 106b plasmid using PEI reagent and incubated for 24 hours in a CO_2_ incubator. After 6 hours of the incubation, the medium was replaced with fresh media and further incubated for 24hours. Following this, the cells transfected with plasmids having fluorescent tags were observed under fluorescent microscope to check the efficiency of transfection

### Isolation of mitochondria

The mitochondria were isolated from the transfected cells, using hypotonic buffer, where the cells were allowed to swell in the buffer for 10 minutes and then break open the cells to release the mitochondria. The cell suspension was then centrifuged at 1300g to remove the cell debris, followed by centrifugation at 12000g to get the mitochondrial pellet. The mitochondrial pellet was suspended in the mitochondrial resuspension buffer.

### Sonication

The mitochondrial pellet was mixed with Lysis buffer and sonicated for 2 minutes at 70% amplitude with 15 sec ON and 30 sec OFF cycle on 4°C. The solution obtained, was centrifuged at 12000g for 10 minutes. The supernatant was collected and protein estimation was done followed by sample preparation for SDS PAGE.

### Protein estimation

Protein level of mitochondria was estimated by Bradford method (Bradford etal,1976). To achieve this, 10μl of sample and 90μl of bradford reagent (50 mg Coomasie Brilliant Blue-G250 in 25ml ethanol and 50ml of phosphoric acid made upto 100ml with water) was added in triplicates in 96 well plate and the absorbance was taken at 595nm by multimode plate reader. The concentration of protein was calculated from the standard plot to BSA with concentration range from 10μg-100μg.

### SDS-PAGE

Protein sample was prepared by mixing of 6x SDS loading dye and boiling it at 90°C for 10 minutes in water bath. The sample was immediately kept on ice and briefly centrifuged before loading on SDS-PAGE gel. The electrophoresis was carried out by using Bio-Rad electrophoresis unit. The protein samples were run through the stacking gel at 80V for 15 minutes and through the resolving gel at 100V at room temperature until the dye reached the end of the gel.

### Western blot analysis

The purity of the mitochondrial pellet was checked by western blot using mitochondria specific antibody (VDAC). Also, the mitochondrial pellet was checked for the nuclear and cytoplasmic contaminants using Histone H3 antibody for Nucleus and Hexokinase HK3 antibody for the cytoplasm.

### RNA Isolation

RNA was isolated from the mitochondrial pellet as well as from the total cell using trizole reagent. Following this, the concentration of the RNA was checked by using nano drop.

### Polyadenylation of RNA

Poly A tail was added to the RNA by Poly A Polymerase enzyme, using manufacturers protocol. This reaction set up was incubated at 37°C for 30 minutes followed by heat inactivation for 5 minutes at 65°C.

### cDNA synthesis

The polyadenylated RNA were used for the synthesis of miRNA 106b specific cDNA by Kang method. Apart from this, total RNA was used to synthesize the cDNA for checking the expression of mitochondrial genes.

### Real Time PCR (qRT PCR)

Quantative real time PCR was performed to check the expression pattern of the microRNA 106b and other mitochondrial genes in mitochondria before and after over expression of miR 106b.

### mRNA stability assay

To elucidate the targeting of mitochondrial genes by miR 106b, mRNA stability assay was performed. The cells were transfected with miR 106b using PEI method. 24 hours post transfection the cells were treated with actinomycine D at 0, 1, 3, 6 and 12 hours. The samples were collected at each time point for gene expression study.

### Oxygraph analysis

To check the phenotypic effects of the down regulation of the mitochondrial genes by miR 106b, the oxygraph analysis was performed, where the oxygen consumption level was checked in the miR 106b over expressed samples and compared with the control samples. In brief, the cells were grown in a 6 well plate and transfected with the candidate microRNAs using PEI method. After 48 hours of incubation at 37°C, the cells were trypsinized and the cell pellet was resuspended in respiration buffer. Later, 1ml of the cellular suspension was added to oxymeter and oxygen intake reading was recorded for 10 minutes. The readings were used to plot the graph to represent the oxygen consumption by the mitochondria.

### Exosome isolation

The exosomes were isolated from the miR 106b over expressing cells using PEG method. To achieve this the cells were transfected with miR 106b and media was replaced with serum free medium. 48 hours post transfection; the spent media was collected and centrifuged at 2000g for 1/2 hour, to remove cell debris. Following this, the cells were mixed with PEG solution in 1:2 ratios and incubated at 4 degree C overnight. Finally, the solution was centrifuged at 12000g for 1 hour. The exosomal pellet was dissolved in PBS and protein estimation was done.

### Cell migration Assay

Cancer cells have the metastatic properties, where they move from its origin to another place and form a secondary tumor. To mimic this in-vitro we perform the cell migration assay with an objective to check, if the exosomes from miR 106b over expressing cancer cells could induce cellular migration of normal Hepatic cells. A scratch was made in the WRL monolayer and treated with exosomes isolated from miR 106b over expressing cancer cells. The cells were allowed to fill the gap formed by the scratch for 48 hours and the images were taken 0, 24, 48 hours respectively and quantified using ImageJ.

### Soft Agar Colony formation assay

To elucidate the role of miR 106b in inducing carcinogenic properties in normal heaptic cells, the colony formation assay was performed where the WRL cells were suspended in the low melting agarose and treated with the exosomes. The cells were incubated at 37°C for 28 days and allowed to form the colonies. The colonies were stained with Coomasie brilliant blue and images were taken from 20 different locations and quantification was done by using ImageJ.

### In-Silico analysis

miR 106b target prediction analysis and scoring for mitochondrial genes was done using five algorithms i.e. MirBase, miRanda, Target Scan, miRDB and MicroRNA.org. Five highest scored mitochondrial genes (ND6, ATP6, Cyto-B, Cox1, ND4L) were selected for target validation.

### Statistical analysis

All the data in the study were expressed as the mean with the standard error mean of at least three experiments, each done in triplicates. SPSS 11.0 software was used for analysis of statistical significance of difference by Duncan’s One way Analysis of Variance (ANOVA). A value of P<0.05 was considered significant.

## Results

### Cellular and mitochondrial levels of miR 106b in HepG2 and WRL cells

First we estimated the cellular and mitochondrial levels of miR 106B in HepG2 cells by keeping WRL cell as control, by RT-PCR analysis. The results revealed that, the level of miR 106B was found to be 2 fold higher in HepG2 cells, when compared with its expression in WRL cells. Next, we checked the expression pattern of miR 106B in the mitochondria of normal & hepatic cancer cells and the results suggested that the abundance of miR 106B was 10 fold lower in the mitochondria of the HepG2 cells when compared with the control cells (WRL).

**Figure 1:**
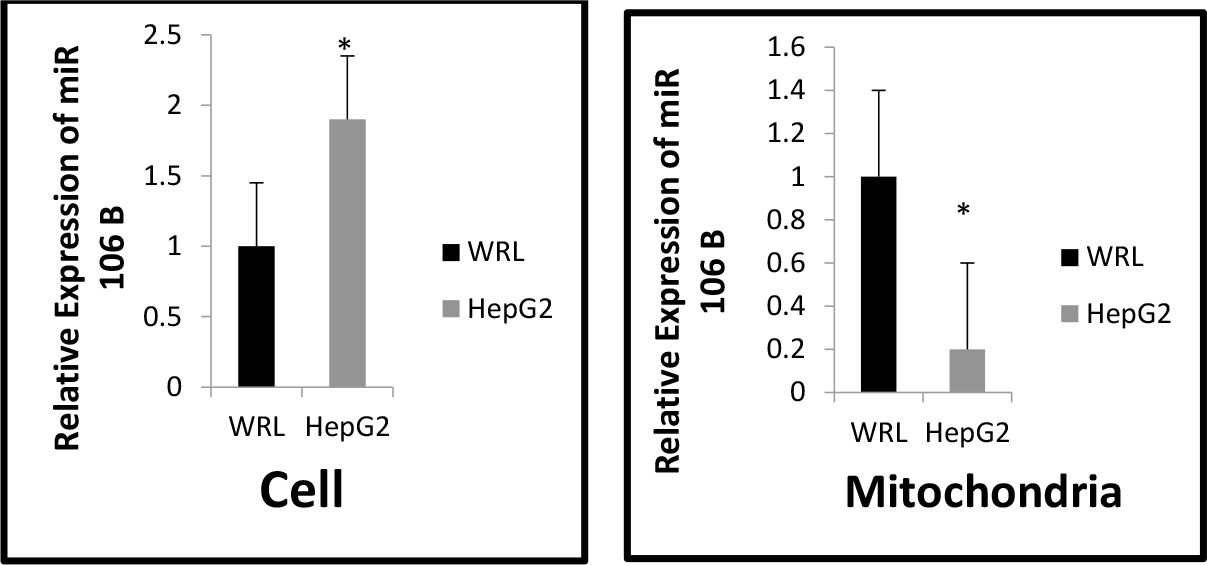
Differential levels of miR 106b in total cell and mitochondria. RT PCR analysis was performed to check the expression pattern of miR 106b in mitochondrial and cellular fraction of HepG2 cells, keeping WRL as control. The results suggested that expression of miR 106b was significantly low in the mitochondria of HepG2 cells. A.) Relative expression levels of miR 106b in the cells B.) Relative levels of miR 106b in the mitochondria. Results presented are average of three experiments ± SEM each done at least in triplicate, p< 0.05. *Statistically significant when compared to control.

### MicroRNA 106b gets targeted to the mitochondria in a selective manner

We found that miR 106b was significantly low in the mitochondria, so next we over expressed it in HepG2 cell line to check if it gets targeted to the mitochondria. The RT-PCR results indicated a 60 fold increase in the levels of miR106b in the mitochondria and 80 fold increase in the cytoplasm, when compared with the control.

**Figure 2:**
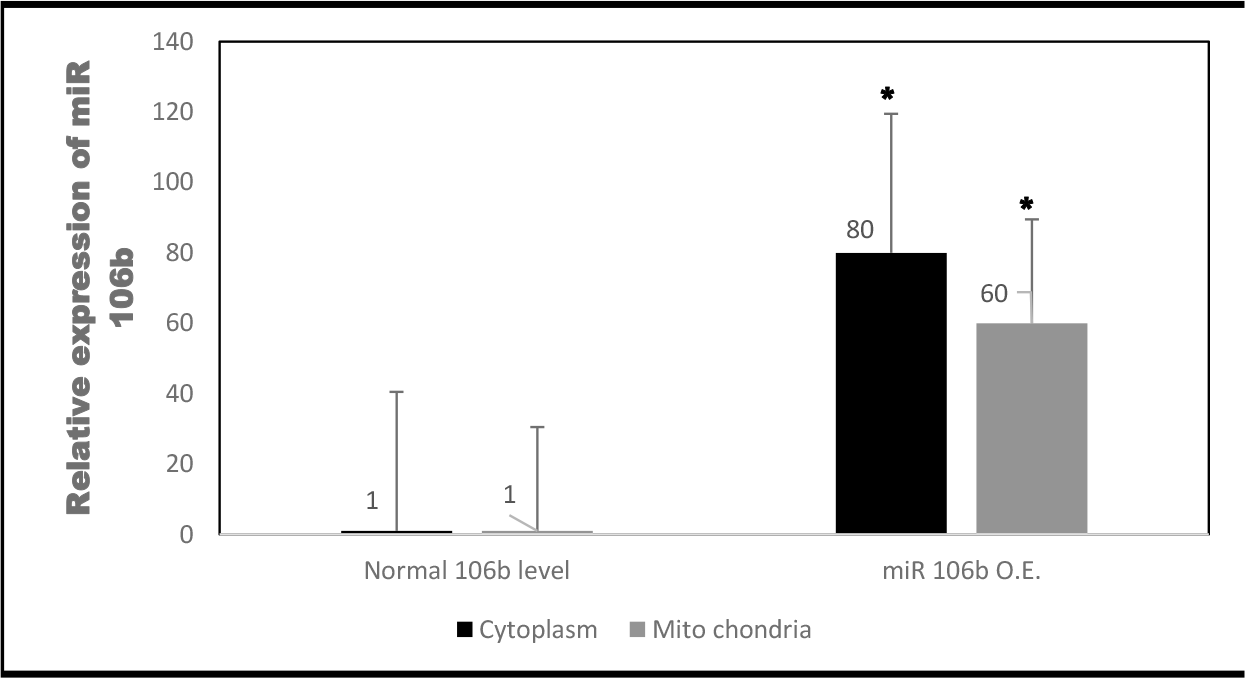
mir 106b gets targeted to the mitochondria in a selective manner. miR 106b was over expressed in HepG2 cell followed by isolation of mitochondria & isolation of RNA from mitochondrial and cytoplasmic fraction. The qRT PCR was performed to check the levels of miR 106b. The results revealed that miR 106b gets targeted to mitochondria in a selective manner with a 60 fold increase in mitochondria and 80 fold increase in cytoplasm. Results presented are average of three experiments ± SEM each done at least in triplicate, p< 0.05.*Statistically significant when compared to control.

### Mitochondrial target genes get significantly down regulated with miR 106b over expression

The target prediction and scoring analysis of miRNA 106b revealed that it has target on important mitochondrial genes. So, after confirmation of the potential localization of miR 106b in the mitochondria, next we checked the impact of this higher levels of miR106b on the expression pattern of their target genes in the mitochondria. To check this, we performed target validation study and the results indicated that expression level of all the target genes of miR106b were significantly lowered, confirming the in-silico target prediction analysis. The level of Cyto B was found to be about 5 folds lowered in the mitochondria of hepatic cancer cells (HepG2) when compared with the non cancerous hepatic cells (WRL). The levels of Cox1, ND4L and ND6 were also found lower, with 1.6, 1.5, and 1.2 folds respectively.

**Figure 3:**
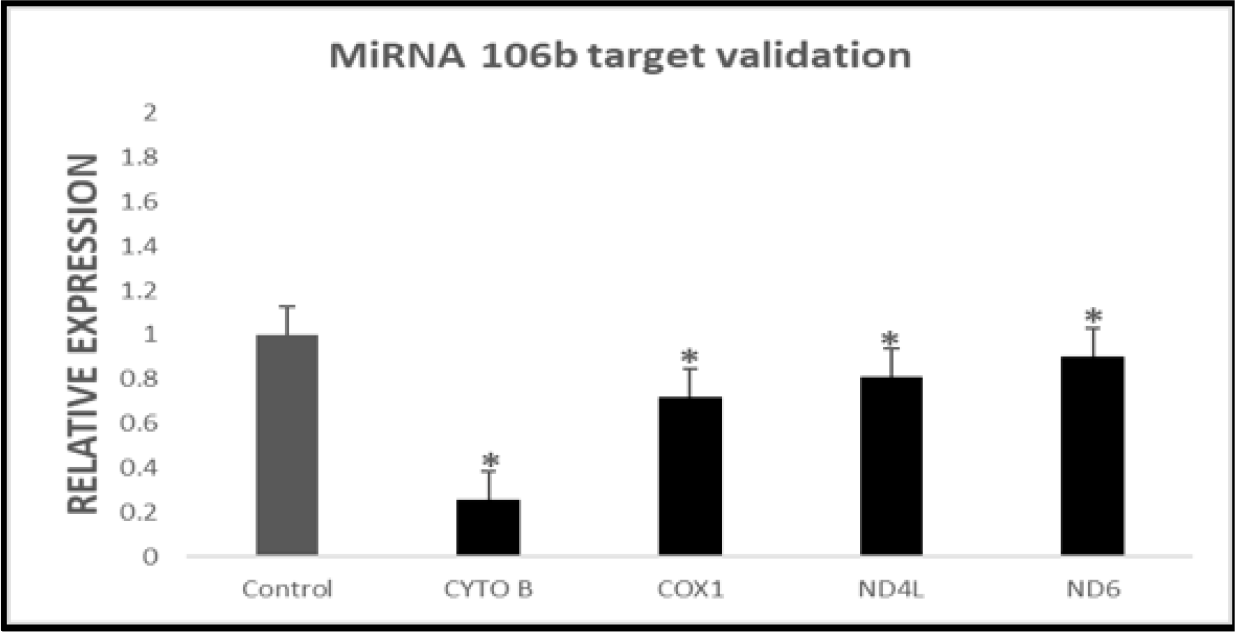
miR 106b down regulates all of its target genes. miRNA 106b was overexpressed in HepG2 cells and the mitochondria isolated. RT-PCR analysis was performed to check the impact of increased miR 106b level on the expression pattern of target mitochondrial genes. The result revealed that the expression of all the mitochondrial target genes of miR 106b went significantly down. Results presented are average of three experiments ± SEM each done at least in triplicate, p<0.05.*Statistically significant when compared to control.

### 3’ UTR analysis and mRNA stability assay confirmed the targeting of mitochondrial genes by miR 106B

The target validation study revealed that the expression levels of all the candidate mitochondrial genes were significantly down regulated by miR 106B. Next, mRNA stability assay was carried out to check the direct targeting of mitochondrial genes by miR 106B and the results revealed that the levels of Cyto B mRNA, was significantly reduced under condition of miR 106b over expression when compared with the levels in mock transfected cells. A consistent lowering pattern was not observed for the levels of CoX1, ND4L and ND6 mRNA.

The results therefore, suggested that Cyto B was effectively targeted by miR106b and falls in line with the 3’ UTR analysis, which revealed that 3’ UTR of Cyto B harbor putative miR106b binding site, whereas the targetting of other mitochondrial genes was not significant.

**Figure 4:**
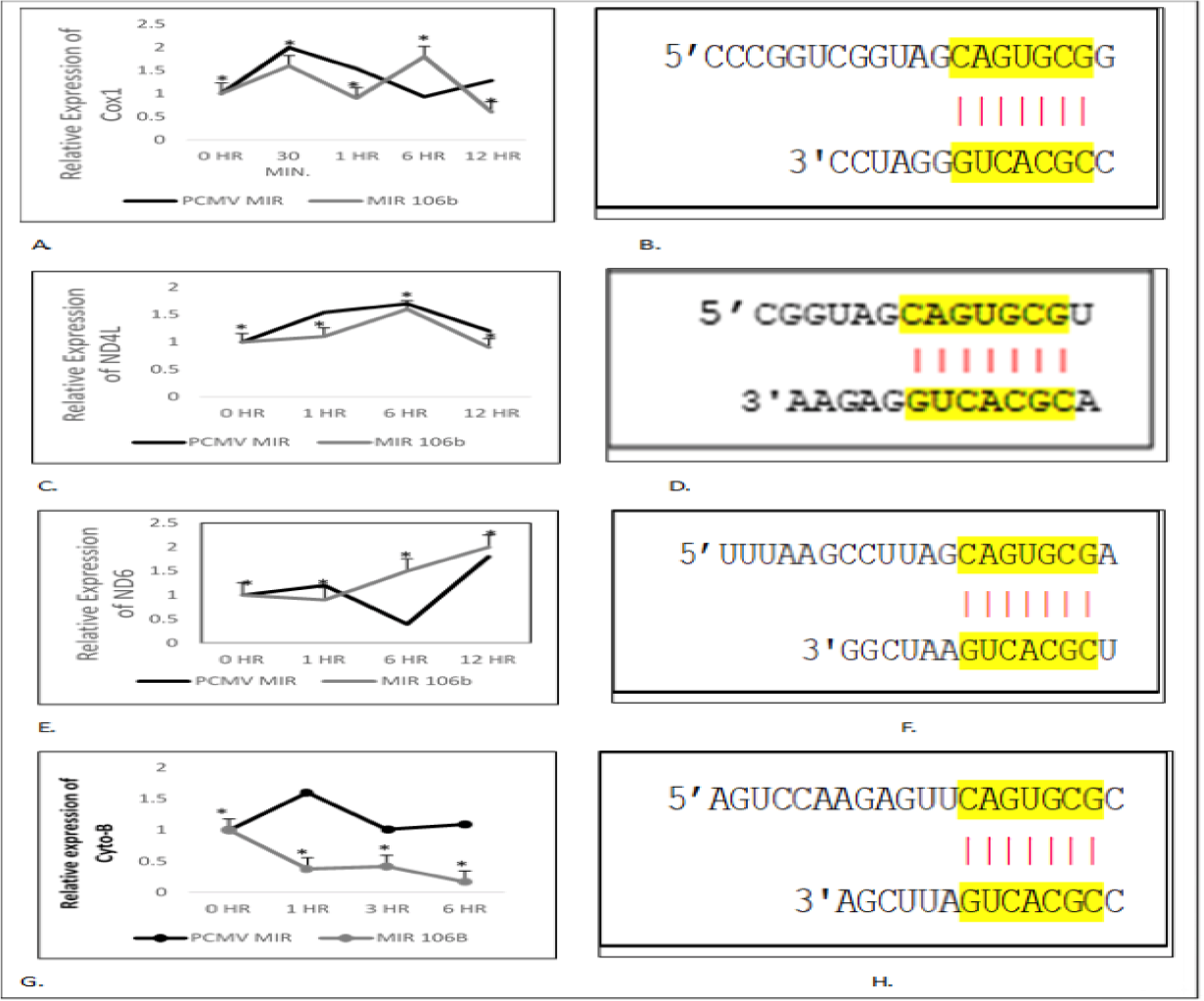
3’ UTR analysis and mRNA stability assay revealed the direct targeting of Cyto-B by miR 106b. miR 106b was overexpressed in HepG2 cells and 24 hours post transfection, the cells were treated with actinomycin D and RNA samples were collected at 0, 1, 6 and 12 hours respectively. Following this, RT-PCR analysis was performed to check the levels of target mRNA. Results revealed that levels of Cyto-B mRNA went significantly down with time suggesting its direct targeting by miR106b. The expression pattern of Cox1, ND6 and ND4L also was altered but not significant. A.) Relative expression of Cox 1 B.) 3’ UTR binding sequence of Cox1 C.)Relative expression of ND4L D.) 3’ UTR binding sequence ND4L E.) Relative expression of ND6 F.) 3’ UTR binding sequence G.) Relative expression of Cyto-B. H.) 3’ UTR binding sequence of Cyto B. Results presented are average of three experiments ± SEM each done at least in triplicate, P<0.05. *Statistically significant when compared to control.

### Mitochondrial oxygen consumption significantly drops upon miR 106b over expression

mRNA stability assay results revealed that miR 106b effectively target Cyto B, which play very important role in the ETC and any alteration in the expression pattern of this gene could significantly alter the functioning of the mitochondrial machinery. To check the phenotypic impact of this down regulation on the functioning of the mitochondria, we performed the oxygraph analysis by checking the levels of the oxygen consumption by the cells. The result of the oxygraph analysis revealed that the levels of oxygen consumption by the cells went significantly down in miR 106b over expressing cells when compared with the control cells, suggesting its role in dysregulating the mitochondrial machinery.

**Figure 5:**
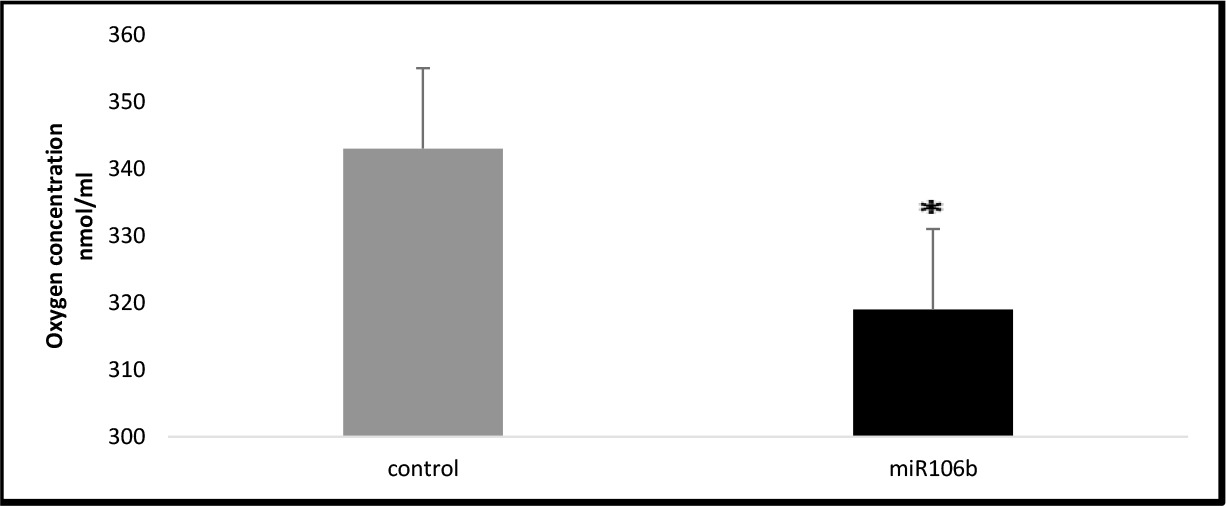
Levels of oxygen consumption by the cells went significantly down, when miR 106b was over expressed. miR 106b was over expressed in HepG2 cell line and 24 hour post transfection, oxygraph analysis was performed. The results revealed a decrease in O_2_ consumption by the mitochondria of miR 106b over expressing cells when compared with the mock transfected cell. Results presented are average of three experiments ± SEM each done at least in triplicate, p< 0.05.*Statistically significant when compared to control.

### Exosomes from miR 106b over expressing cancer cells escalate the rate of cell migration and stimulate the colony formation in cancerous and non-cancerous hepatic cells

The results till now revealed that miR 106b gets targeted to the mitochondria and down regulate the expression of Cyto B gene, important in ETC, and it resulted in the reduced oxygen consumption by the HepG2 cells, possibly due to the altered mitochondrial metabolism. So, we next checked the role of this altered mitochondrial metabolism in oncogenesis.

The results of cell migration assay revealed an increase in the rate of cellular migration when treated with exosomes isolated from the miRNA 106 B over expressing cells, compared to the cells treated with the exosomes isolated from the mock transfected cells and the pattern was found to be persistent in both, cancerous as well as non-cancerous hepatic cell lines.

Along with cell migration assay, we also performed the colony formation assay to check the carcinogenic ability of miR 106b in hepatic cells. Here, we treated the WRL & HepG2 colonies with the exosomes isolated from miR 106b over expressing or mock transfected cells. Colony formation assay result, shown an increase in the number and size of the colonies when treated with the exosomes enriched with miR106B when compared with the colonies treated with exosomes isolated from the mock transfected cells.

**Figure 6:**
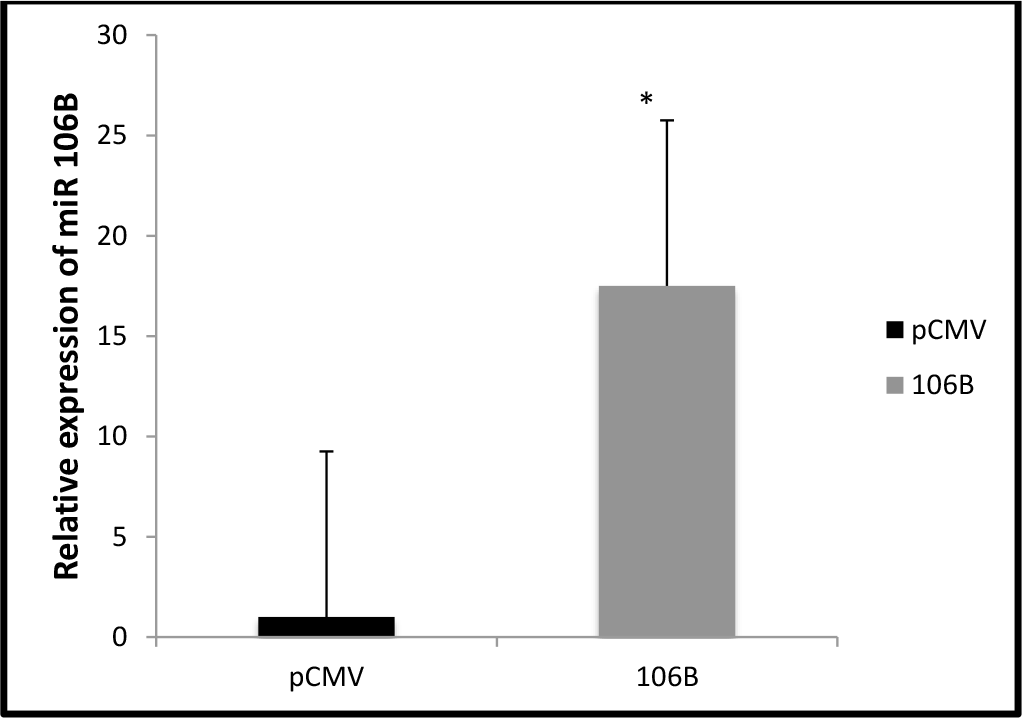
Exosomes isolated from miR 106b over expressing cells were enriched with miR 106b. Real time PCR analysis was performed to check the enrichment of miR106b in the exosomes isolated from miR 106b over expressing cells and compared with the exosomes isolated from the control cells. The results revealed that exosomes isolated from microRNA 106b over expressing cells, shown 17.5 fold higher level when compared with control. Results presented are average of three experiments ± SEM each done at least in triplicate, p<0.05. *Statistically significant when compared to control.

**Figure 7:**
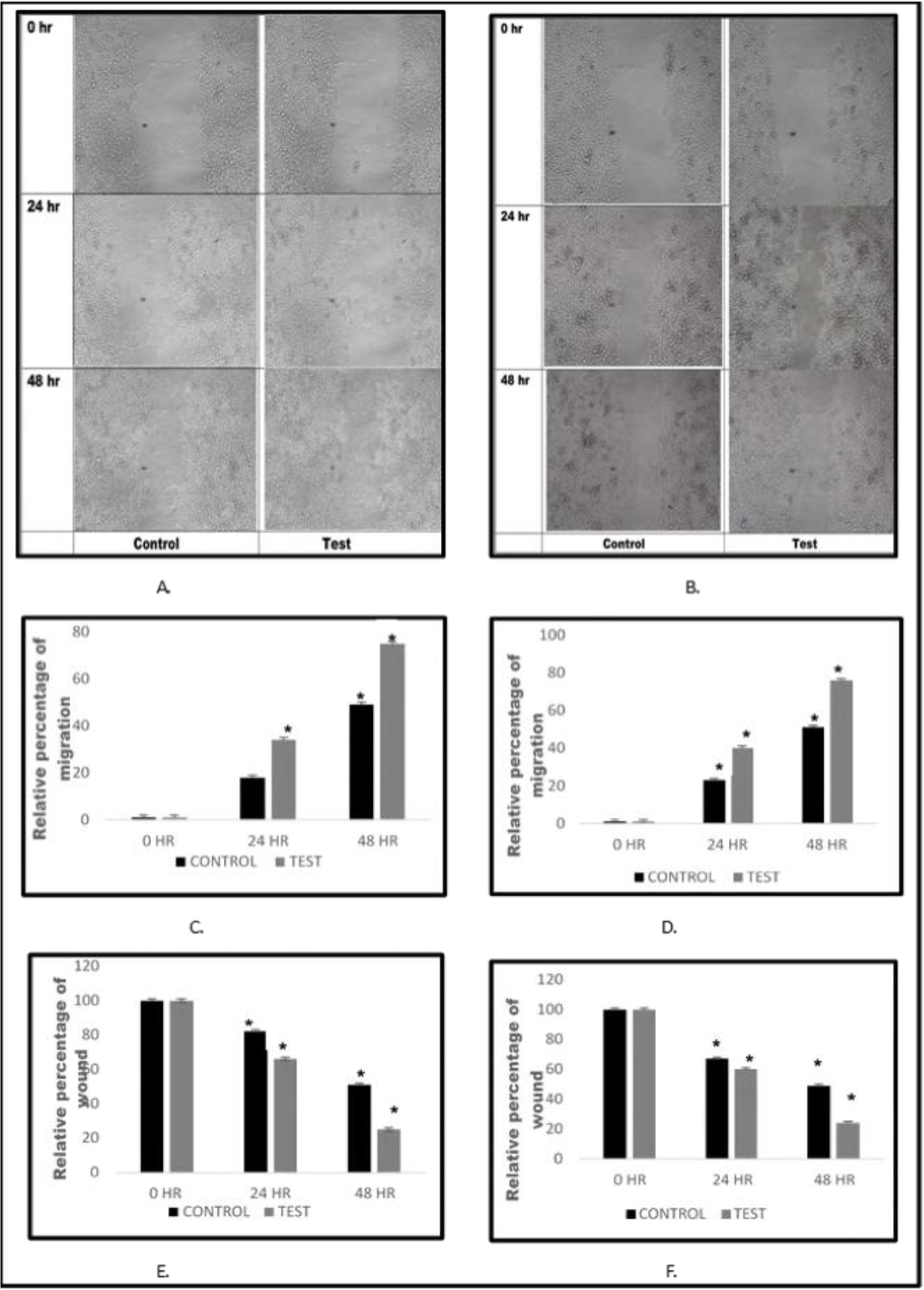
miR 106b escalates the cellular migration when treated with exosomes isolated from microRNA 106b over expressing cells. Cells were grown as a monolayer and a scratch was made, followed by exosome treatment. The microphotographs were taken at 0, 24, and 48 hours. The distance/gap covered by the cells with time was estimated by image-J software. The result revealed that the rate of the cellular migration got escalated when treated with exosomes enriched with miR 106b. A.) Microphotograph of cell migration pattern (WRL) with respect to time B.) Microphotograph of cell migration pattern (HepG2) with respect to time C.) Relative percentage of migration by WRL cells at 0, 24 & 48 hours D.) Relative percentage of migration by HepG2 cells at 0, 24 & 48 hours E.) Relative percentage of wound closure (WRL cells), at 0, 24 & 48 hours. F.) Relative percentage of wound closure (HepG2 cells), at 0, 24 & 48 hours. Results presented are average of three experiments ± SEM each done at least in triplicate, p<0.05. *Statistically significant when compared to control.

**Figure 6.8:**
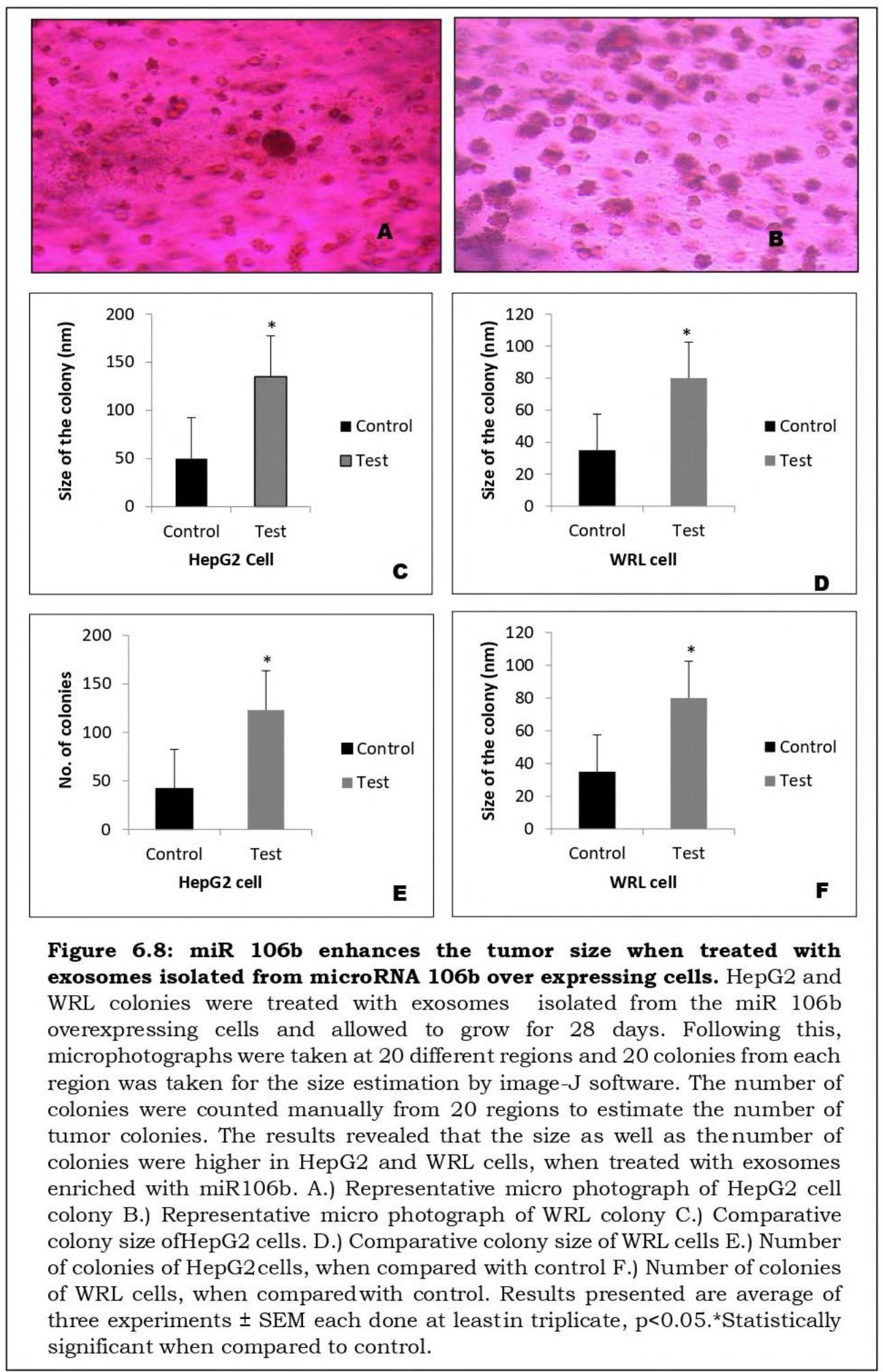
miR 106b enhances the tumor size when treated with exosomes isolated from microRNA 106b over expressing cells. HepG2 and WRL colonies were treated with exosomes isolated from the miR 106b overexpressing cells and allowed to grow for 28 days. Following this, microphotographs were taken at 20 different regions and 20 colonies from each region was taken for the size estimation by image-J software. The number of colonies were counted manually from 20 regions to estimate the number of tumor colonies. The results revealed that the size as well as the number of colonies were higher in HepG2 and WRL cells, when treated with exosomes enriched with miR106b. A.) Representative micro photograph of HepG2 cell colony B.) Representative micro photograph of WRL colony C.) Comparative colony size ofHepG2 cells. D.) Comparative colony size ofWRL cells E.) Number of colonies of HepG2cells, when compared with control F.) Number of colonies of WRL cells, when compared with control. Results presented are average of three experiments ± SEM each done at leastin triplicate, p<0.05.*Statistically significant when compared to control.

## Discussion

Cancer is the second most fatal disease around the globe (Nagai H et. al, 2017). It is a disease of unregulated cellular proliferation (Krieghoff-Henning E et. al, 2017). The process involving the initiation and progression of cancer is called carcinogenesis and the agents inducing cancer are known as carcinogens (Stewart B et. al, 2019). Cancer cells have ability to bypass all the antigrowth signals (Ravi S et. al, 2022), which is acquired by the multiple mutations in the genes regulating the cell growth and proliferation (Basu A et. al, 2018). The mutations in such important genes either leads to their altered function or changes their expression pattern i.e. high or low (Sinkala M et. al, 2023).

Apart from the mutations, the change in expression pattern of the genes involved in carcinogenesis, could also be due to the involvement of microRNAs (Hagan J et. al, 2007). MicroRNAs are a class of non-coding RNAs of 22-25 nucleotides capable of regulating the expression pattern of a variety of genes important in normal functioning of the cells (Brien J et. al, 2018). MicroRNAs play crucial role in carcinogenesis as they target both the classes of genes involved in cancer i.e. Oncogenes and Tumor suppresor genes (TSG) (Fasoulakis Z et. al, 2020).

Various microRNAs have been reported to be the key players in carcinogenesis. The miRNAs invloved in cancer are classified in two groups, i.e. oncomiRs and tumor suppressor miR (Svoronos A et. al, 2016). OncomiRs basically target the tumor suppressor genes and regulate their function, whereas the tumor suppressor miRNAs regulate the expression of oncogenes (Zhang B et. al, 2007). MicroRNA 106b is located on 7th chromosome at 7q22.1 (Hudson R et. al, 2013) and it has been reported to be involved in various types of cancers, by regulating the expression level of genes involved in cellular proliferation, invasion and metastasis (Yang F et. al, 2022, Sagar SK, 2021).

Several studies revealed the involvement of microRNAs in the regulation of mitochondrial genes and hence, altering its normal functioning (Rodrigues S et. al, 2020, Zhang G et. al, 2021, Borralho P et. al, 2015). Since mitochondria is an important cell organelle, which supplies energy to the cells for their normal functioning, any alteration would change the cellular metabolism and hinder the normal functioning (Giulivi C et. al, 2023). Altered mitochondrial energetics has been reported as one of the important hallmark of cancer (Hanahan D, 2022).

The microRNAs which regulate expression levels of the mitochondrial genes could either be of mitochondrial origin or get localized to mitochondria from the cytoplasm (Parmasivam A et. al, 2020). This class of microRNAs, involved in regulation of expression levels of mitochondrial genes are called mitomiRs (Patel D et. al, 2023). The main objective of this study was to check the role of miR 106B in modulation of the mitochondrial metabolism and impact of this reprogramming in oncogenesis. Since, miR 106B is a nuclear coded miRNA, so first we checked its levels in the mitochondria by qRT-PCR and found that its expression was significantly low in the mitochondria. Several studies suggest that nuclear coded micro RNAs gets targeted to the mitochondria (Das S et. al, 2012). So, next we overexpressed miR 106B in hepG2 cells and the RT-PCR results revealed that the, miR 106b got enriched in the mitochondria by 60 folds, in a selective manner.

As our in-silico analysis revealed that miR 106b have target on the important mitochondria genes crucial in electron transport chain, so next we performed the target validation analysis to check if miR 106b could alter the expression pattern of target mitochondrial genes. The qRT-PCR results revealed that miR 106b lowered the expression level of all its target genes, most significantly Cyto-B. To further prove the targeting of mitochondrial genes by miR106b, we performed mRNA stability assay and the results revealed the targeting of Cyto-B by miR106b and falls in line with the 3’ UTR analysis that revealed that 3’ UTR of Cyto B harbor putative miR 106b binding sites. mRNA stability assay also revealed that miR 106b target Cox1, ND6 and ND4L to a lower extent.

Since, Cyto-B is a crucial gene involved in the complex 3 of electron transport chain, its down regulation would reprogram the mitochondrial mechanism and its effect could be seen phenotypically. To check the phenotypic effects, we performed the oxygraph analysis to monitor the oxygen consumption by mitochondria of miR106b over expressing cells. The oxygraph analysis showed a decrease in the oxygen consumption by the mitochondria of miR 106b over expressing HepG2 cell when compared with the control cells, suggesting its role in dysregulating the mitochondrial energetics by targeting Cyto-B, which is crucial for electron transport chain.

Next, we performed cell migration and colony formation assay to check if over expression of miR 106b can induce carcinogenesis in hepatic cells. The results of colony formation assay and cell migration assay, revealed an increase in the size of the colony when treated with the exosomes enriched with miR106b and cell migration assay also revealed an increase in the rate of cellular migration when treated with these exosomes when compared with the cells treated with exosomes isolated from the mock transfected cells. These results therefore, suggest that miR106b possess carcinogenic effect and it possibly involve the modulation of mitochondrial metabolism.

## Acknowledgment

We acknowledge Indian Council of Medical Research, Ministry of Health, Govt. of India for the financial assistance in form SRF and Kerala state council for Science Technology & Environment, Govt. of Kerala for fellowship in the form of JRF and SRF to Mr. Ashutosh K. Maurya. We also acknowledge Central University of Kerala for providing all the necessary facilities to carry out this research work.

## Author Contributions

The authors confirm contribution to the paper as follows: Study conception and design: VBSK, Bioinformatics and wet lab work: AKM. Cloning of miR 106b: LE. All authors reviewed the results and approved the final version of the manuscript.

## SUPPLEMENTARY DATA

